# Age-dependent dysregulation of locus coeruleus firing in a transgenic rat model of Alzheimer’s disease

**DOI:** 10.1101/2022.11.15.516661

**Authors:** Michael A. Kelberman, Jacki M. Rorabaugh, Claire R. Anderson, Alexia Marriott, Seth D. DePuy, Kurt Rasmussen, Katharine E. McCann, Jay M. Weiss, David Weinshenker

**Affiliations:** Department of Human Genetics, Emory University, Atlanta, GA, USA; Eli Lilly, Indianapolis, IN, USA; Department of Psychiatry and Behavioral Sciences, Emory University, Atlanta, GA, USA

**Author notes:** JM Rorabaugh now works at Teva Pharmaceuticals, Denver, CO, USA. CR Anderson now works at Eurofins Lancaster Laboratories, Lancaster, PA, USA. SD DePuy now works at Premier Consulting, Morrisville, NC, USA. K Rasmussen now works at Delix Therapeutics, Boston, MA, USA.

**Keywords:** locus coeruleus, tau, electrophysiology, Alzheimer’s disease, TgF344-AD, aging

## Abstract

Accumulation of hyperphosphorylated tau in the locus coeruleus (LC) is a ubiquitous feature of prodromal Alzheimer’s disease (AD), and LC neurons degenerate as AD progresses. Tau-mediated LC dysfunction may contribute to early neuropsychiatric symptoms, while loss of LC integrity is associated with conversion to cognitive impairment. Hyperphosphorylated tau alters firing rates in other brain regions, but its effects on LC neurons have not been described. The purpose of this study was to characterize changes in firing properties of LC neurons when they are the only cells containing hyperphosphorylated tau, as well as later in disease when β-amyloid (Aβ) and tau pathology is abundant in the forebrain. Single unit LC activity was recorded from anesthetized wild-type (WT) and TgF344-AD rats, which carry the APP/PS1 transgene. Similar to human AD, these rats develop hyperphosphorylated tau in the LC (at 6 months) prior to Aβ or tau pathology in forebrain regions (at 12-15 months). At baseline, LC neurons from TgF344-AD rats were hypoactive at both ages compared to WT littermates, but showed elevated spontaneous bursting properties, particularly in younger animals. Differences in footshock-evoked LC firing depended on age, with 6-month TgF344-AD rats demonstrating aspects of hyperactivity, and aged transgenic rats showing hypoactivity relative to WT. Tau-induced alterations in LC firing rates may contribute to the pathophysiology of AD, with early hyperactivity associated with prodromal symptoms, followed by hypoactivity contributing to cognitive impairment. These results support further investigation into disease stage-dependent noradrenergic interventions for AD.

**Highlights:**

- Recorded locus coeruleus (LC) neurons in a rat model of Alzheimer’s disease (AD)
- TgF344-AD rats develop early endogenous LC tau pathology akin to human AD
- 6- and 15-month TgF344-AD rats had reduced tonic LC firing
- LC neurons from 6-month TgF344-AD rats were hyperactive in response to footshock
- LC neuron dysfunction may contribute to AD symptoms

## 1. Introduction

Accumulation of hyperphosphorylated tau within subcortical nuclei and subsequent dysfunction of these neurons is a nearly ubiquitous feature along Alzheimer’s disease (AD) progression [1, 2]. A seminal report from Braak and colleagues [3], independently replicated by other groups [4-6], positions the noradrenergic locus coeruleus (LC) as the earliest site of pathological tau deposition, well before cortical β-amyloid (Aβ) plaque accumulation or the onset of diagnostic cognitive deficits. During prodromal phases of AD, non-cognitive symptoms consistent with noradrenergic hyperactivity, including sleep disturbances, agitation, and anxiety, emerge coincident with the appearance of hyperphosphorylated tau in the LC [7–11]. Cerebrospinal fluid norepinephrine (NE) levels and its turnover are elevated in early AD [12–15], and a recent neuroimaging study demonstrated that higher LC signal on a neuromelanin-sensitive MRI was predictive of neuropsychiatric symptom severity in AD patients [16], further supporting the theory of LC hyperactivity during initial stages of disease. At the same time, numerous studies have linked the deterioration of LC integrity to cognitive and structural decline in aging and AD [17–22], suggestive of reduced LC-NE transmission during later stages of the disease.

While neurochemical, neuropathological, and behavioral results are consistent with disease stage-specific alterations in LC activity, direct evidence for changes in LC firing is mostly lacking. Aβ pathology often induces neural hyperactivity [23], including in the LC [24], whereas tau pathology typically induces neuronal hypoactivity [25]. However, there are other reports of tau-mediated hyperactivity [26–28], suggesting region and/or cell-type specific effects. Given that Aβ only accumulates in the LC during late stages of AD [24, 29], the dysregulation of LC circuits at the level of the cell bodies is likely dominated by the early accumulation of hyperphosphorylated tau. Therefore, understanding the impact of aberrant tau on LC neural activity is critical for determining the neurobiological underpinnings of prodromal AD symptoms and progression to cognitive impairment. This information could then inform rational development of early biomarkers and therapeutic interventions at various disease stages.

The objective of this study was to delineate the effects of AD-like hyperphosphorylated tau on LC firing rates. Though some studies have begun to track LC activity in humans using functional MRI [17], these techniques lack spatial and target specificity for small regions like the LC [30]. We therefore employed the TgF344-AD rat model, which expresses mutant human amyloid precursor protein and presenilin-1 (APP/PS1) that cause autosomal dominant, early-onset AD [31]. This model possesses several benefits for our study. These rats demonstrate many of the same behavioral phenotypes that are observed in AD that are influenced by LC activity. These include early anxiety-like behaviors followed later by cognitive impairment that can be reversed by LC activation [9, 10, 31, 32]. In addition, TgF344-AD rats, unlike their APP/PS1 transgenic mouse counterparts, develop endogenous tau pathology that first appears in the LC [32]. This tau deposition is coincident with the appearance of non-cognitive behavioral abnormalities but prior to tau or Aβ pathology elsewhere in the brain, reminiscent of human disease progression. In the current study, we recorded single unit LC activity from anesthetized TgF344-AD rats and wild-type (WT) littermates at baseline and in response to footshock at 6 months, an age when anxiety-like behavior emerges and hyperphosphorylated tau in the LC is the only detectable AD-like neuropathology, as well as 15 months, when brain-wide tau and Aβ pathology are evident in combination with deficits in learning and memory.

## 2. Methods

### 2.1 Animals

This study used a total of 42 TgF344-AD rats and WT littermates on a Fischer background aged 6 or 15 months. TgF344-AD rats were hemizygous for the *APPsw/PS1ΔE9* transgene that contains mutations causative of autosomal dominant early-onset AD [31]. Rats were housed in groups of 2-3 on a 12-h light/dark cycle (lights on at 7:00 am) with food and water available *ad libitum.*

All experiments were conducted in accordance with the Institutional Animal Care and Use Committee at Emory University. Male and female rats were assigned to sex-balanced experimental groups, given the lack of prominent sex differences noted in this strain [9, 31, 33].

### 2.2 Surgery

At two months of age, rats to be used for electrophysiology underwent stereotaxic surgery. Rats were anesthetized with 5% isoflurane and maintained at 2% throughout surgery. Prior to incision, rats were given ketoprofen (5 mg/kg, s.c.). An AAV9-PRSx8-mCherry-WPRE-rBG virus was infused bilaterally targeting the LC (AP: −3.8 mm, ML: +/- 1.2 mm, DV: −7.0 mm from lambda with the head tilted 15 degrees downward). The injection syringe was left in place for 5 min following the infusion prior to being moved dorsally 1 mm and waiting an additional 2 min to ensure diffusion of virus at the site of injection. This virus was to be a control for a planned experiment that we did not end up pursuing, and nothing further was done with it in this study. Electrophysiology recordings were performed approximately 4 or 13 months following surgery.

### 2.3 Electrophysiology

At 6 or 15 months, rats were anesthetized with chloral hydrate (400 mg/kg, i.p.) and secured in a stereotaxic frame. An incision was made to expose the skull, which was leveled based on measurements made at bregma and lambda. A 15° head tilt was employed to avoid the sagittal sinus, and burr holes were drilled over the approximate location of the LC (AP: 3.8-4.0 mm, ML: 0.9-1.3 mm from lambda). 16-channel silicone probes (V1×16-Poly2-10mm-50s-177-V16_100-50, NeuroNexus; Ann Arbor, MI) were connected to a u-series Cereplex headstage (Blackrock Neurotech; Salt Lake City, UT). A 16-channel Cereplex Direct System was used to acquire digitized signals with a 250 Hz-5 kHz bandpass filter and 10 kS/s sampling rate. LC units were identified based on field standard criteria, including stereotaxic coordinates, location adjacent to the mesencephalic trigeminal nucleus (Me5), biphasic response to footpinch/footshock, and reduction/cessation of spontaneous activity following injection of the selective α2-adrenergic receptor agonist clonidine (0.1 mg/kg i.p.). Each recording began with a 5-min baseline period, which was immediately followed by 10 applications of a contralateral footpinch for LC verification, each separated by 10 s, as described previously [34]. Afterwards, 0.5 ms 10 mA footshocks (each separated by 10 s for 5.5 min) were applied to the contralateral hindpaw, followed by the same pattern using 5 ms 10 mA footshocks. Footshocks were delivered by an ISO-Flex stimulus isolator and controlled by a Master-8 (A.M.P. Instruments; Jerusalem, Israel). LC spikes were manually sorted using Blackrock Offline Spike Sorting Software.

Electrophysiology analysis was performed in NeuroExplorer v5 (Nex Technologies; Colorado Springs, CO) or Matlab (R2019a; Mathworks; Natick, MA). To ensure that single units were being analyzed, neurons with >2% of recorded spikes within a predefined 3 ms refractory period were eliminated [35]. The 5-min baseline recording served as the basal firing rate and to calculate the interspike interval for each single unit. Spontaneous bursting properties (number of bursts, percentage of spikes within a burst, burst duration, spikes per burst, interspike interval within a burst, burst rate, and interburst interval) of LC neurons during baseline recordings were also quantified. Spontaneous bursts were defined as two spikes with an interspike interval of <0.08 s and terminated with an interspike interval >0.16s, as previously described [36, 37]. Footshock was used to ascertain changes to sensory evoked LC activity. LC response to footshock was divided into three response categories: immediate (0-60 ms), intermediate (60-100 ms), and late (200-400 ms). These periods were based on a previous report demonstrating that altering the length (0.5, 2, and 5 ms) of footshock could elicit both a standard immediate and long latency LC response [38].

### 2.4 Tissue Preparation and Immunohistochemistry

Following completion of electrophysiological recordings, rats were overdosed with isoflurane and perfused with potassium phosphate-buffered saline followed by 4% paraformaldehyde (PFA). Brains were removed and stored in 4% PFA overnight then transferred to 30% sucrose until sectioning. Brain sections containing the LC were sliced at 30 um and either dry mounted or stored in cryoprotectant until being processing for immunohistochemistry.

Dry mounted sections were counterstained with neutral red. Slides were submerged in increasing concentrations of ethanol (70%, 95%, and 100%) for 1 min each, followed by 10 min in neutral red, dunked 3-5 times in increasing concentrations of ethanol (70%, 95%, and 100%), and finally in xylene for 2 min. Slides were coverslipped with permount and imaged on a Keyence BZ-X700 microscope (KEYENCE; Osaka, Japan).

To confirm that α2-adrenergic receptors *(Adra2a)* are specific to noradrenergic neurons in the LC region and thus the only cells that could respond to clonidine during our recordings, we performed fluorescent *in situ* hybridization using the RNAscope Multiplex Fluorescent V2 Assay (Advanced Cell Diagnostics, Newark, CA, USA) on brainstem sections containing the LC. One 6-month WT male was lightly anesthetized, and the brain was quickly removed and flash frozen in isopentane on dry ice. The brain was stored at −80C until sectioning on a cryostat at 16μm. The RNAscope assay was performed following the manufacturer’s protocol, multiplexing tyrosine hydroxylase (*Th*) and *Adra2a.* Images were taken in a 1μm pitch z-stack (10um total) on a Keyence BZ-X700 microscope at 20x and 40x. A representative 2x section was also captured to visualize most of the coronal section containing the LC.

### 2.5 Statistical Analysis

Data are presented as raincloud plots consisting of a density plot, box-and-whisker plot, and individual data points that were created in R using modified code from open-source platforms and from [39, 40]. All code is available upon reasonable request. Statistical analysis was performed using GraphPad Prism (v. 9.2.0; San Diego, CA). A two-way ANOVA was used to identify main effects of age, genotype, or their interaction. When applicable, post-hoc tests were performed across genotypes within an age group using the Holm-Sidak correction.

## 3. Results

### 3.1 LC Neural Recording Verification

We used neuroanatomical, sensory, electrophysiological, and pharmacological methods to verify that recordings were from bona fide LC neurons. LC units were located medial to Me5, which was identified by jaw deflection **(Figure 1A)**. Putative LC units displayed a canonical biphasic response to footpinch/footshock, and as previously reported, a subset of these neurons demonstrated a late phase (200-400 ms) response to 5 ms footshock that was not observed in response to 0.5 ms footshock **(Figure 1B)**. LC cells, unlike other adjacent nuclei, express high levels of *Adra2a* and are inhibited by the α2-adrenergic receptor agonist clonidine [41] **(Figure 1C, D)**. Finally, we verified that electrode tracks were in the LC post-recording using a neutral red counterstain **(Figure 1E)**.

**Figure 1.**
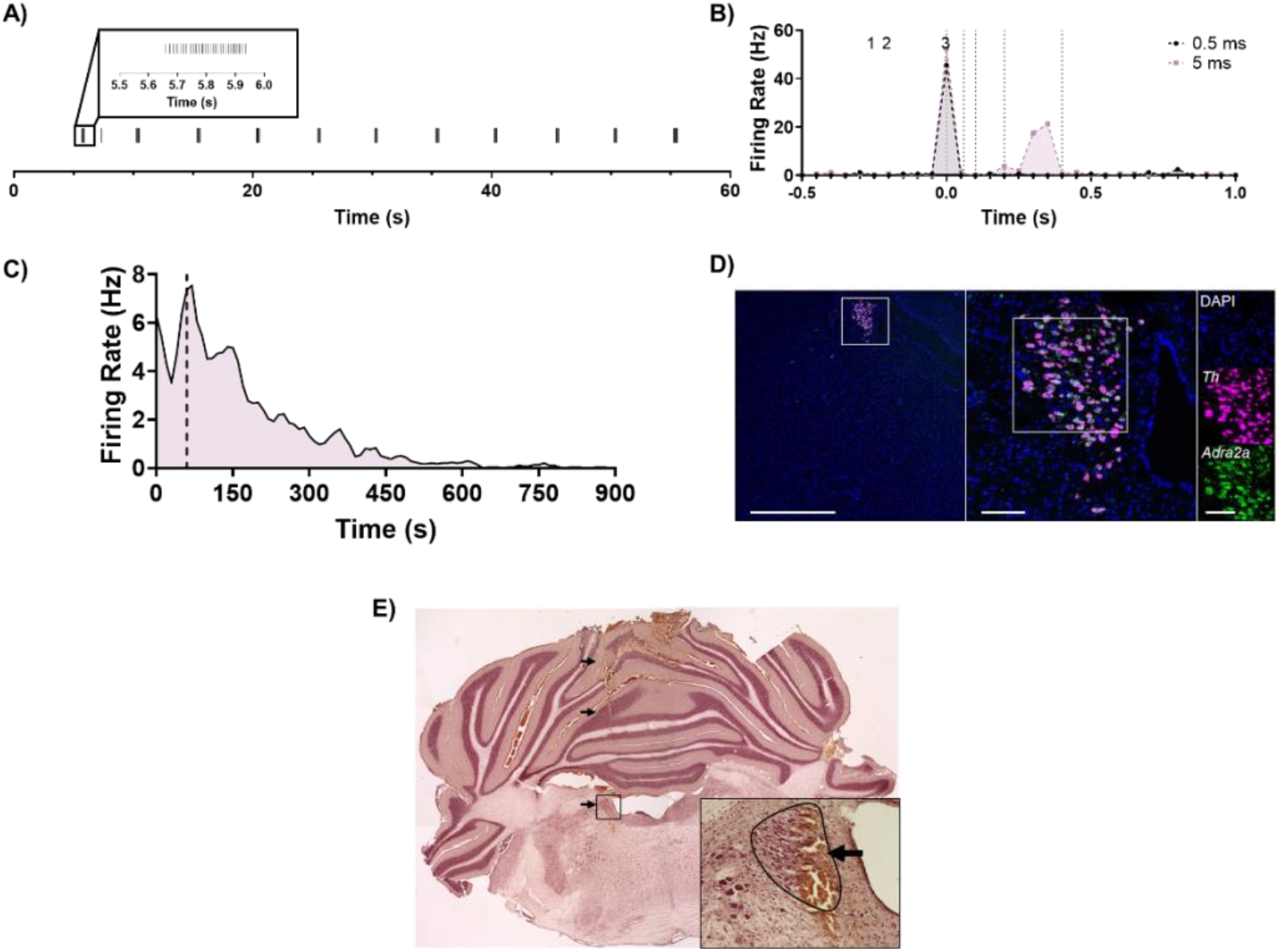
Multimodal confirmation of single unit recordings from LC neurons. A) The approximate depth of the LC was estimated by locating Me5 via jaw deflection every 5 s during 1-min long recording. B) Putative LC units displayed a biphasic response to footshock (gray, 0.5 ms; purple, 5 ms; 0.05 ms bins around footshock at time 0 s). Unit responses were also quantified between 0-60 ms (1), 60-100 ms (2), and 200-400 ms (3) following footshock. C) Noradrenergic LC neuron activity is inhibited by the α2-adrenergic receptor clonidine (0.1 mg/kg i.p.; dashed line), and D) RNAscope performed at the level of the LC confirmed that expression of α2-adrenergic receptors (green) is limited to TH-expressing (magenta) LC neurons in this part of the brain. Scale bars: 1 mm at 2x, 100 μm at 20x, and 100 μm at 40x magnification. E) Localization of electrode tracks (arrows) in the LC core using neutral red counterstaining. Main image stitched from four images at 2x magnification. Inset taken at 20x magnification with the LC outlined in black and the electrode track marked by an arrow.

### 3.2 Alteration of Pacemaker-like LC Firing in TgF344-AD Rats

LC neurons fire with regular pacemaker activity between 0.5-2 Hz under normal conditions that maintains baseline levels of arousal, attention, and noradrenergic tone [42, 43]. To assess tau pathology- and age-induced changes in tonic firing, we recorded periods of spontaneous activity from 75-120 isolated LC neurons per group (N=8-11 animals). There was a significant main effect of genotype (F_1, 385_ = 4.35, p = 0.04) on baseline firing rates, such that LC neurons of TgF344-AD rats were less active than those from WT littermates **(Figure 2A)**. There was no effect of age (F_1, 385_ = 0.1 1, p = 0.74) or an age x genotype interaction (F_1, 385_ = 0.78, p = 0.38). We also quantified interspike interval for units with two or more spikes (N=74-119 neurons/group), defined as the time between successive action potentials **(Figure 2B)**. There was a trend towards a main effect of genotype (F_1, 379_ = 3.55, p = 0.06), where TgF344-AD LC neurons had a shorter time between successive spikes, which was surprising given that tonic firing was lower in the TgF344-AD rats. However, LC neurons can also transiently fire in brief bursts, even in the absence of external stimuli [37, 44–47]. Therefore, we quantified the spontaneous bursting properties of LC neurons from TgF344-AD and WT rats. Of the neurons exhibiting spontaneous bursts (N=48-88 neurons/group), there was a main effect of genotype on firing rate within a burst (F_1, 267_ = 6.57, p = 0.01) and on interspike interval within a burst (F_1, 267_ = 8.02, p < 0.01). During spontaneous bursts, firing rates were higher in TgF344-AD rats and interspike intervals were lower. There were no alterations in other aspects of spontaneous bursts (**Table 1**). Overall, these changes indicate lower basal firing rates but elevated bursting properties of LC neurons in TgF344-AD rats.

**Figure 2.**
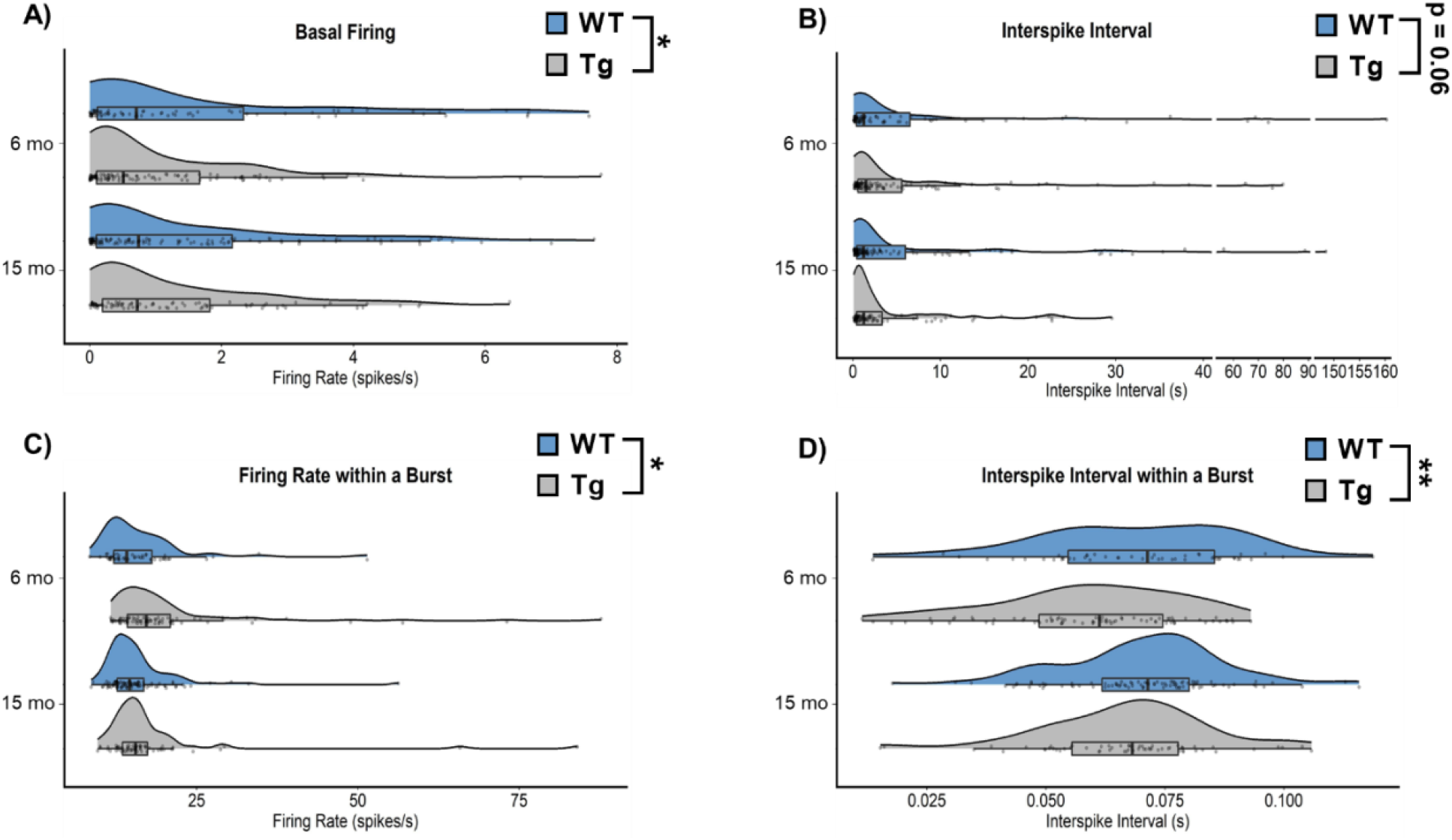
Basal firing properties of LC neurons from 6- and 15-month TgF344-AD rats and WT littermates. A) Baseline firing was lower in TgF344-AD rats at both ages. B) There was a trend towards decreased interspike interval in TgF344-AD rats. C) Firing rate during spontaneous bursts was higher in TgF344-AD rats. C) Interspike intervals within a burst were lower in TgF344-AD rats, and there was an additional effect of age where younger rats had lower interspike intervals. N=74-120 neurons/group (N=8-11 animals/group) for A and B. N=48-88 neurons/group (N=8-11 animals/group) for C and D. *p<0.05, **p<0.01.

**Table 1.** Spontaneous bursting properties of LC neurons.

| **Measure** | **Main Effect** | **F Statistic** | **p-value** | **Summary Values [Mean +/- SEM (N)]** |  |  |
| --- | --- | --- | --- | --- | --- | --- |
| **Number of Bursts** | **Age** | $F_{1,385} = 0.002712$ | 0.9585 | | **WT** | **Tg** |
| | **Genotype** | $F_{1,385} = 3.258$ | 0.0719 | **6 month** | 26.89 ± 6.00 (75) | 16.43 ± 3.29 (107) |
| | **Interaction** | $F_{1,385} = 0.6039$ | 0.4376 | **15 month** | 25.53 ± 3.58 (120) | 19.37 ± 3.45 (87) |
| **Percent of Spikes in a Burst** | **Age** | $F_{1,385} = 1.649$ | 0.1999 | | **WT** | **Tg** |
| | **Genotype** | $F_{1,385} = 0.1737$ | 0.6771 | **6 month** | 18.41 ± 2.75 (75) | 19.64 ± 2.37 (107) |
| | **Interaction** | $F_{1,385} = 0.8379$ | 0.3606 | **15 month** | 23.84 ± 2.23 (120) | 20.55 ± 2.44 (87) |
| **Burst Duration** | **Age** | $F_{1,267} = 0.5341$ | 0.4655 | | **WT** | **Tg** |
| | **Genotype** | $F_{1,267} = 2.652$ | 0.1046 | **6 month** | 0.36 ± 0.07 (48) | 0.32 ± 0.04 (75) |
| | **Interaction** | $F_{1,267} = 0.5149$ | 0.4737 | **15 month** | 0.44 ± 0.06 (88) | 0.32 ± 0.02 (60) |
| **Spikes Per Burst** | **Age** | $F_{1,267} = 0.9097$ | 0.3411 | | **WT** | **Tg** |
| | **Genotype** | $F_{1,267} = 0.06193$ | 0.8037 | **6 month** | 7.46 ± 1.41 (48) | 8.61 ± 1.56 (75) |
| | **Interaction** | $F_{1,267} = 1.356$ | 0.2453 | **15 month** | 7.72 ± 1.08 (88) | 5.94 ± 0.34 (60) |
| **Interburst Interval** | **Age** | $F_{1,226} = 0.01437$ | 0.9047 | | **WT** | **Tg** |
| | **Genotype** | $F_{1,226} = 2.222$ | 0.1375 | **6 month** | 16.27 ± 3.67 (37) | 37.83 ± 8.13 (60) |
| | **Interaction** | $F_{1,226} = 3.516$ | 0.0621 | **15 month** | 27.51 ± 5.21 (75) | 2505 ± 5.42 (58) |

### 3.3 Dysregulated LC Response to Footshock in TgF344-AD Rats

LC neurons fire in transient bursts in awake animals to a variety of salient stimuli, including novelty, tones, pain, and stress [48–50]. We characterized the responsiveness of LC neurons 0-60, 60-100, and 200-400 ms following footshock (10 mA, 0.5 or 5 ms), which maintains its ability to trigger LC bursting under anesthesia, as described [38]. For the immediate response phase (**Figure 3A, 3B**), there was a reduction of LC activity in TgF344-AD rats that was specific to the 0.5 ms footshock (F_1, 433_ = 6.06, p = 0.01). There was a significant age x genotype interaction on LC firing rates in the mid-phase response to both 0.5 ms (F_1, 433_ = 6.26, p = 0.01) and 5 ms (F_1_, 452 = 9.56, p < 0.01) footshock (**Figure 3C, 3D**). LC neurons from TgF344-AD rats were hyperactive at 6 months following the 5 ms footshock (t_452_ = 2.36, p = 0.04), but hypoactive at 15 months following the 0.5 (t_433_ = 2.16, p = 0.03) and 5 ms footshock (t_452_ = 2.03, p = 0.04), compared to age-matched WT littermates. We observed no main effects on late phase response to either 0.5 ms or 5 ms footshock (**Figure 3 E & F**).

**Figure 3:**
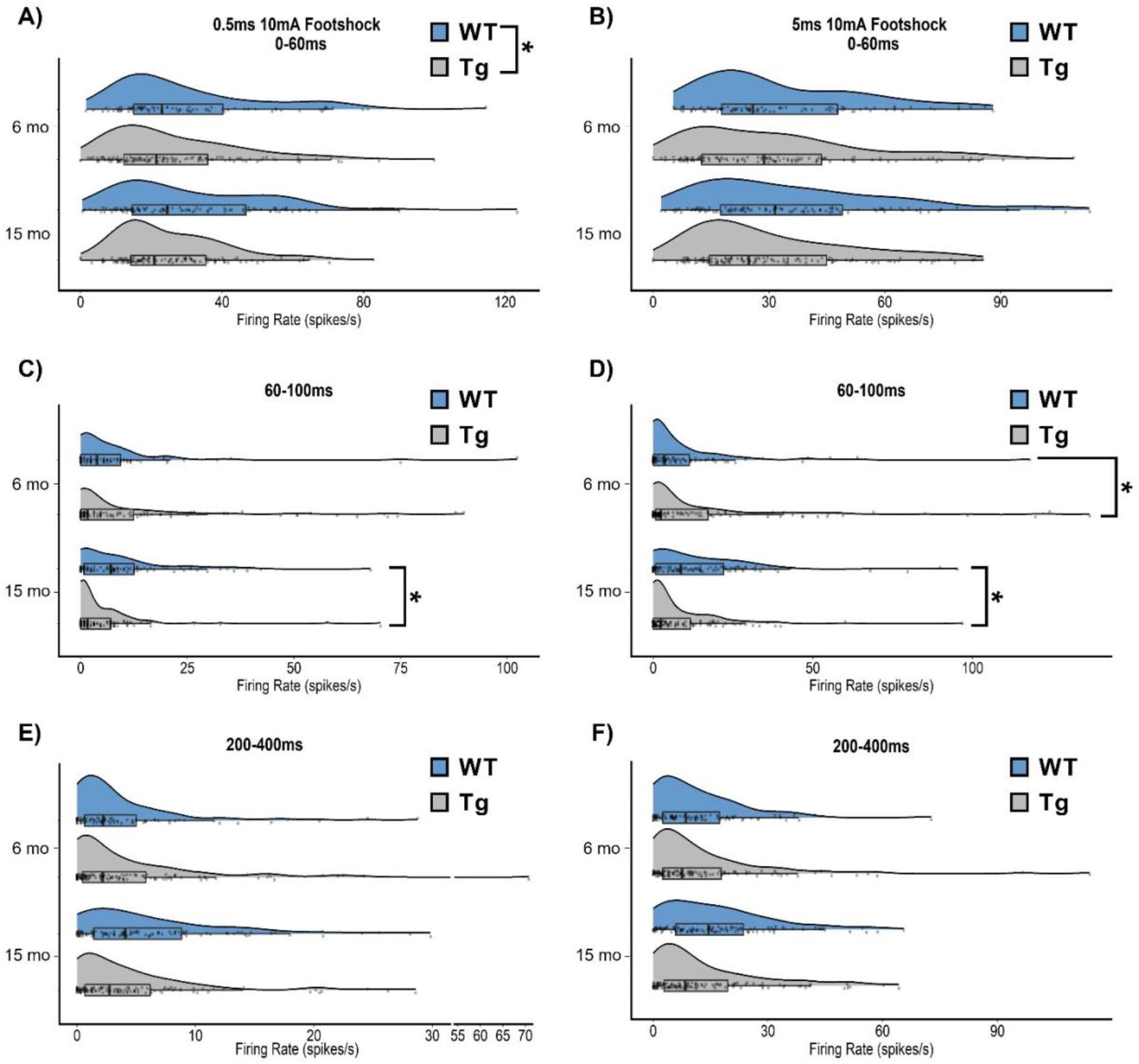
Footshock response of LC neurons from 6- and 15-month TgF344-AD rats and WT littermates. A & B) LC neuron firing rate was lower in TgF344-AD rats during the immediate phase in response to 0.5 ms footshock, but this effect was not seen in response to 5 ms footshock. C & D) There was an age x genotype interaction in the intermediate phase of response to both 0.5 and 5 ms footshock. Post-hoc analysis revealed that young TgF344-AD animals demonstrated hyperactivity in response to 5 ms footshock, while aged TgF344-AD animals showed hypoactivity in response to both 0.5 and 5 ms footshock. E & F) There were no differences in late phase response to footshock. N=84-128 neurons/group (N=8-11 animals/group) for A, C, and E. N=94-126 neurons/group (N=8-11 animals/group) for B, D, and F. *p<0.05.

## 4. Discussion

### 4.1 Overview of changes in LC firing and associations with symptoms of AD

The LC is one of the earliest regions to develop AD-related neuropathology in the form of hyperphosphorylated tau and undergoes frank cell death in mid- to late-stage disease [3–6, 51]. The appearance of hyperphosphorylated tau in the LC is coincident with the emergence of prodromal symptoms of AD such as sleep disturbances, agitation, dysregulated mood, and increased anxiety, which are suggestive of LC hyperactivity [7–11]. Early LC hyperactivity is further supported by increases in cerebrospinal fluid NE levels and turnover [12–15], axonal sprouting and elevated receptor density [52, 53], and adrenergic receptor hypersensitivity [54]. By contrast, late-stage AD is characterized by phenotypes consistent with NE deficiency such as apathy and cognitive impairment that are correlated with loss of LC integrity [21, 55–58]. These data suggest that early hyperphosphorylated tau triggers compensatory mechanisms that increase LC-NE transmission and maintain function, which eventually fail as LC neurons succumb to more advanced pathology [52–54]. The divergent phenotypes across the progression of AD could be explained, at least in part, by changes in LC firing rates.

Prior to our study, only two investigations tracked changes in LC activity using rodent AD models, one with Aβ and one with mutant P301S tau pathology [24, 59]. Mislocalization of GABA receptors associated with soluble Aβ oligomers in the LC led to hyperactivity, whereas effects of P301S tau expression were minimal. There are limitations in these studies that should be noted. In both cases, only one aspect of AD-like pathology was incorporated, the transgenes were driven by ubiquitous promoters, and LC Aβ and P301S tau are more reminiscent of late-stage AD and frontotemporal dementia, respectively. It is therefore unsurprising that we uncovered different dysregulated tonic and phasic LC firing patterns using an AD model that develops both Aβ and tau pathology (**Figure 4**), the latter of which is comprised of endogenous wild-type tau and isolated to the LC at early ages.

**Figure 4:**
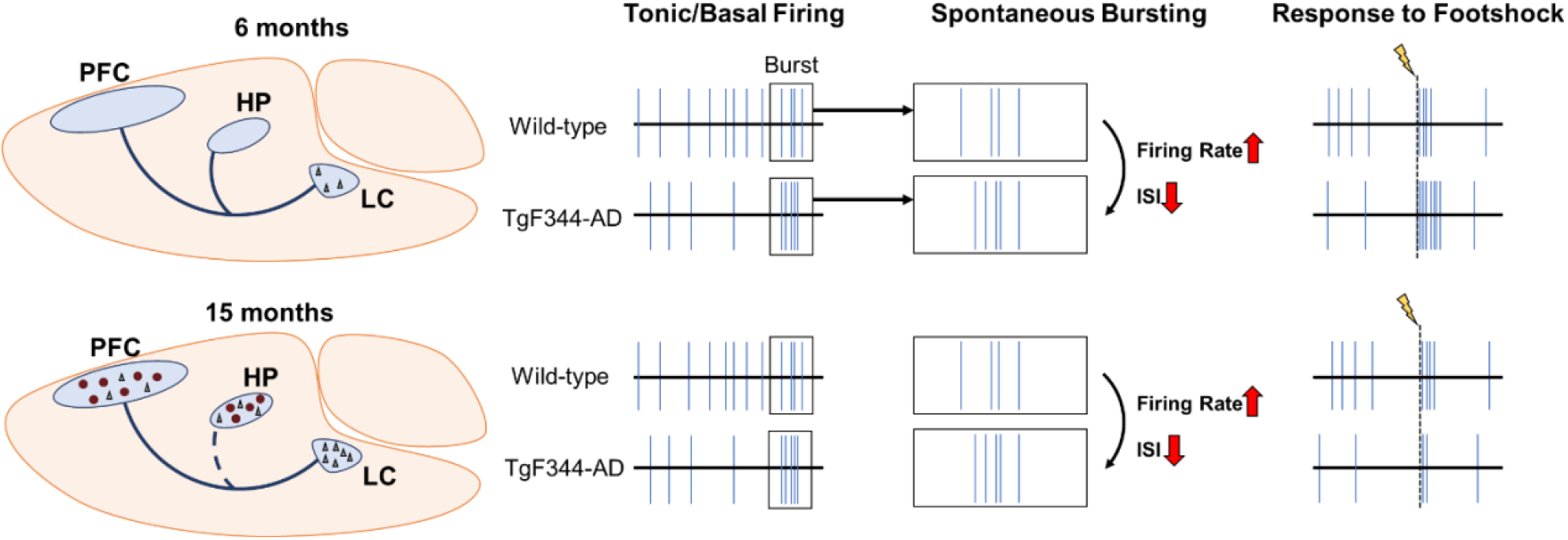
Summary of dysregulated LC firing patterns as a function of age in TgF344-AD rats. At 6 months, the LC is the only brain region that displays AD-like neuropathology in the form of hyperphosphorylated tau (black triangles). LC neurons show tonic hypoactivity, but elevated firing rate and shorter interspike intervals during periods of spontaneous bursts. LC response to footshock (lightning bolt/dashed line) at this age is also elevated in TgF344-AD rats. At 15 months, hyperphosphorylated tau pathology in the LC worsens, and tau and Aβ (red circles) pathology are present throughout the forebrain. There is also evidence of noradrenergic denervation, specifically to the hippocampus (dashed blue line). Similar to 6-month TgF344-AD rats, LC neurons again show tonic hypoactivity, and elevated firing rate and lower interspike intervals during periods of spontaneous bursts. However, in contrast to younger rats, 15-month TgF344-AD rats have an impaired, rather than elevated, footshock response.

In our study, both young and old TgF344-AD animals showed tonic LC hypoactivity but elevated spontaneous bursting properties. Moreover, LC activity in response to footshock stress was elevated in 6-month TgF344-AD rats. Importantly, we and others have reported behavioral abnormalities in young TgF344-AD reminiscent of neuropsychiatric symptoms in prodromal AD [8–11, 60]. At first glance, reduced pacemaker LC firing is inconsistent with evidence that excessive NE transmission contributes to comorbidities seen in prodromal phases of AD [7]. Specifically, elevations in tonic, but not phasic, LC activity are associated with stress and anxiety [61–63]. On the other hand, increased phasic LC activity driven by spontaneous bursting and/or environmental stimuli could promote improper transitions through different stages of arousal [46, 64, 65], resulting in fractured sleep/wake cycles that are common in AD [7, 8]. The importance of distinguishing between different patterns of LC firing across disease stages is highlighted by a recent report that enhancement of phasic LC firing protects LC neurons from deleterious forms of tau, whereas high tonic firing results in AD-associated psychiatric symptoms and worsened neuronal health [66]. Together, these data are consistent with the idea that a combination of homeostatic mechanisms to maintain LC function in the presence of pathology and damage (e.g. altered LC firing, increased NE turnover, elevated adrenergic receptor sensitivity and density, and axonal sprouting) contributes to prodromal behaviors. These homeostatic/compensatory mechanisms ultimately fail in later disease, as indicated by reduced NE tissue levels, noradrenergic denervation, and frank LC neuron loss [9, 32, 54, 55, 67, 68], and our results showing tonic LC hypoactivity and blunted response to footshock in 15-month TgF344-AD rats. This culminates in a loss of LC-NE transmission and cognitive impairment, which our lab has successfully rescued by selectively augmenting tonic LC firing [32, 69, 70].

### 4.2 Potential mechanisms underlying changes in LC firing rates

While the neurobiological mechanisms underlying the changes in LC firing properties we observed remain to be determined, the nature of those changes offers clues. Reductions in tonic firing rates and hypoactivity in aged TgF344-AD rats are unlikely due to cell death because there is no LC neuron loss at this age [32]. However, we know that β-adrenergic receptor function in the dentate gyrus is heightened in young TgF344-AD rats, indicating that adrenergic receptor plasticity exists in this model [54]. In the pons, α2-adrenergic receptors are abundantly expressed within the LC itself (**Figure 1**) and regulate its activity as inhibitory autoreceptors. Changes in the density or sensitivity of these receptors, which have been noted in the human condition with other subtypes of adrenergic receptors [52–54], could alter LC neuron firing properties. Furthermore, elevated phasic activity may result in greater NE release compared to tonic activity [71]. A surplus of NE at the level of LC cell bodies would culminate in augmented inhibition from neighboring LC neurons, also potentially leading to tonic LC hypoactivity.

Besides its intrinsic pacemaker activity, the LC also receives extensive excitatory, inhibitory, and modulatory serotonin (5-HT), hypocretin/orexin, corticotropin releasing hormone) input from various cortical and subcortical regions that could contribute to altered firing rates in TgF344-AD rats [72, 73]. GABAergic inputs arising from the ventrolateral preoptic area or the peri-LC region modulate basal noradrenergic tone and have been mainly studied in the context of arousal [74–76]. In addition, given that the LC and peri-LC GABAergic neurons receive a set of non-overlapping innervations [74], it is possible that changes in firing rates are the result of a dysregulated, multi-synaptic pathway. For example, systemic application of a 5-HT receptor agonist lowers tonic but enhances phasic firing of the LC in a GABA-dependent manner [77]. Similarly, 5-HT dampens sensitivity of LC neurons to glutamatergic inputs, which originate from the cortex, lateral habenula, and paragigantocellularis nucleus [78–82]. Thus, altered serotonergic signaling to the LC represents a singular mechanism that could influence both altered tonic and phasic firing patterns. With respect to footshock-evoked LC activity, the immediate and middle phase responses are also mediated, in part, by excitatory amino acids [81, 82]. Changes in excitatory amino acid transmission, receptor expression, or localization could be responsible for various aspects of dysregulated LC firing. Other neuropeptides, such as hypocretin/orexin and corticotropin releasing factor powerfully activate LC neurons and are both associated with elevations of tonic LC firing and the expression of anxiety-like phenotypes [61–63, 83, 84]. The TgF344-AD rats represent an excellent model to further evaluate AD pathology-associated changes in these systems and their influence on LC activity.

### 4.3 Clinical Implications

We have previously posited that LC/NE-based therapies for AD should be guided by disease stage [7]. The dysregulated/hyperactive NE transmission that is produced by early AD pathology (e.g. hyperphosphorylated tau) and promotes prodromal symptoms may be best alleviated by therapies that decrease LC firing and/or NE signaling, while reduced LC-NE transmission resulting from advanced pathology and deterioration of LC neurons would benefit from LC-NE stimulation. Our current findings largely support this model. At 6-months of age, TgF344-AD rats have hyperphosphorylated tau in the LC but no degeneration and display phenotypes relevant to neuropsychiatric disorders such as anxiety, and they primarily present with LC hyperactivity (increased firing during bursts and in responses to footshock stress). Meanwhile 15-month transgenic animal have increased tau pathology, some loss of LC fibers/terminals, cognitive impairment, and display LC hypoactivity (reduced baseline and footshock-induced firing). Moreover, we previously reported that deficits in reversal learning in 15-month TgF344-AD rats can be rescued by chemogenetic LC stimulation [32], and we predict that drugs that suppress LC activity or antagonism of adrenergic receptors would be effective at alleviating prodromal phenotypes in young TgF344-AD rats [7]. These results provide a foundation for translation to the clinical setting, where promising data already exist. For example, adrenergic antagonists have been reported to reduce agitation/aggression and anxiety in subjects with probable or possible AD [85, 86], and our recent phase II study showed beneficial effects of the NE transporter inhibitor atomoxetine, which increases extracellular NE levels, on AD biomarkers in mild cognitive impairment [87]. Atomoxetine also improves response inhibition in Parkinson’s disease patients, with the greatest benefits observed in those with lower LC integrity [88]. Whether this is also true in AD has not been determined but will be important for guiding the use of personalized noradrenergic-based interventions.

## 5. Conclusions

We have identified disease stage-dependent changes in LC firing patterns in a rat model of AD that accumulates hyperphosphorylated tau in the LC prior to forebrain pathology. These data further support the notion that early LC hyperactivity and late LC hypoactivity contribute to prodromal symptoms and cognitive/memory impairments of AD, respectively. These insights should prove useful for developing noradrenergic-based therapeutics for AD.

## Funding and Acknowledgements

This work was supported by funding from the National Institutes of Aging (AG062581 to DW, AG069502 to MAK), the National Institute of Neurological Disorders and Stroke (NS96050 to MAK), and the Eli Lilly Innovation Fellowship Award (JMR and CRA).

We would like to thank A Korukonda for acquiring the image for Figure 1 E and CH West for helping establish the electrophysiological methods used in this study.

This research project was supported in part by the Viral Vector Core of the Emory Center for Neurodegenerative Disease Core Facilities and the Vector Core of the University of Pennsylvania.

## CRediT authorship contribution statement

**Michael A. Kelberman:** Conceptualization, Methodology, Software, Validation, Formal analysis, Investigation, Resources, Data Curation, Writing – Original Draft, Writing – Review & Editing, Visualization, Funding Acquisition. **Jacki M. Rorabaugh:** Conceptualization, Methodology, Investigation, Writing – Review & Editing, Funding Acquisition. **Claire R. Anderson:** Conceptualization, Methodology, Investigation, Writing – Review & Editing, Funding Acquisition. **Alexia Marriott:** Investigation, Writing – Review & Editing. **Seth D. DePuy:** Methodology, Writing – Review & Editing. **Kurt Rasmussen:** Methodology, Writing – Review & Editing. **Katharine E. McCann:** Writing – Review & Editing, Software, Visualization. **Jay M. Wiess:** Conceptualization, Methodology, Writing – Review & Editing, Resources, Supervision, Funding Acquisition. **David Weinshenker:** Conceptualization, Methodology, Validation, Formal analysis, Resources, Writing – Original Draft, Writing – Review & Editing, Visualization, Supervision, Funding Acquisition.

